# Ecological flexibility and selectivity in mixed-species flock participation in birds

**DOI:** 10.1101/2022.11.08.515689

**Authors:** Laura Vander Meiden, Ian R. Hoppe, Daizaburo Shizuka, Allison E. Johnson

## Abstract

Mixed-species groups are hypothesized to allow animals to minimize competitive interactions and maximize facilitative interactions. Individuals’ participation in mixed-species groups may reduce rates of competition and increase the social information available about predators or food availability. Behavioral plasticity may further increase these benefits as plastic species alter their rates of niche overlap with group mates. We investigate two axes of behavioral plasticity that may modulate how species interact with group mates in mixed-species groups—flexibility and selectivity. Specifically, we assess avian species’ patterns of selective preferences for participation in flocks of certain strata and whether behavioral flexibility in foraging strata corresponds with the foraging strata of flock mates. All species in our study maintained or increased their foraging strata overlap with flock mates, supporting the hypothesis that facilitation plays an important role in flock formation. Notably, the methods that species used varied: some species moved closer to flock mates via flexibly matching their flock mates’ behavior, some showed selectivity for flocks of certain stratums, and others did both. Ultimately, we show that species balance facilitative and competitive interactions with flock mates via multiple methods and that consideration of behavioral plasticity is integral to understanding the nuances of mixed-species flock interactions.

## Introduction

In species interactions, individuals must balance the negative impacts of competitive interactions with the potential benefits of facilitative interactions. Mixed-species groups are a useful model system for looking at the interplay of these competitive and facilitative interactions, due to the high rate of interactions between species that vary in their rates of niche overlap (Goodale et al. 2010). Defined as two or more species foraging and moving together though a landscape, mixed-species groups (Morse 1970) are formed by many taxa including birds, fish, primates, ungulates and cetaceans (Greenberg 2000; Lukoschek and McCormick 2000; Stensland et al. 2003) and are found throughout the world in a variety of ecosystems. These groups can be ephemeral or persist for long periods of time (Goodale et al. 2017), and membership can be dynamic, with individuals joining and leaving at different points in time and space, or stable (Martínez and Gomez 2013; Johnson et al. 2018). Much of the literature on mixed-species groups has focused on the identification of nuclear species—that is, species that contribute to the facilitation and cohesion of flocks (Moynihan 1962)—and the benefits these species may provide to satellite or follower species. However, focusing exclusively on the interactions between nuclear and satellite species is limiting, since group membership is shaped by the balance of competition and facilitation among all participating species. Thus, each mixed-species group contains a complicated web of both facilitative and competitive interactions, and different species may navigate this web in different ways.

The dynamics of mixed-species groups are particularly well-studied in birds, where mixed-species flocks commonly form across different ecosystems (Sridhar et al. 2012). Classically, two non-mutually exclusive hypotheses have been proposed to explain why birds participate in mixed-species flocks. First, the reduced competition hypothesis posits that foraging with heterospecific individuals may reduce competitive interactions relative to foraging with conspecifics who compete for the same resources. This competition hypothesis predicts that species will reduce competition by preferentially associating with species of different niches or by changing their foraging behavior to increase niche differentiation in the presence of similarly foraging species (Morse 1977). Evidence from the neotropics has long supported this theory, finding that congeners (Graves and Gotelli 1993) or species from similar foraging guilds (Colorado and Rodewald 2015) were often found in a checkerboard distribution across the landscape rather than in the same flocks.

Second, the more recently conceived ecological facilitation hypothesis posits that heterospecific flock mates may provide higher quality information or information of a different scope about food resources and predator presence than that provided by conspecific individuals, producing facilitative interactions that benefit flocking individuals (Sridhar and Guttal 2018). This hypothesis predicts that flock participants will maximize the utility of benefits provided by preferentially associating with species *more* similar to themselves than expected by chance or by adjusting their behavior to increase niche similarity to those around them. Evidence has been found to support a prominent role for such facilitation in flock formation at both global and local scales, finding that similarities between species’ body mass and diet drive co-occurrence patterns (Zou et al. 2011; Sridhar et al. 2012).

Thus, current evidence for these hypotheses is mixed; we can see signatures of both reduced competition and ecological facilitation playing roles in structuring mixed-species flocks. However, the majority of the studies described above have been focused on how species’ flock selectivity, i.e. species presence in certain flock compositions and not others, correlates with similarity in static traits such as body size, foraging guild, and phylogenetic relatedness. Using these static traits as a proxy for species’ niche similarity fails to account for the role of behavioral plasticity in foraging behavior in flocks. Behavioral plasticity is a key aspect of mixed-species associations, and studies have illustrated that mixed-species flocks can act as vehicles of niche expansion and plasticity. When participating in mixed-species flocks in comparison with conspecific flocks, members may shift their foraging to more risky substrates and strata (Zou et al. 2011), shift their behavior towards higher rates of insectivory (Valburg 1992), and even exhibit an increased likelihood of discovering novel food sources (Freeburg et al. 2017). However, the bulk of research on plasticity in mixed-species flocks has focused on how species change their behavior in and out of mixed-species flocks or specifically in the presence or absence of a particular nuclear species (Dolby and Grubb 1998; Hsieh and Chen 2011), with little work done on how the behavior of a focal species may vary given the behavior of its flock mates as a whole.

Here, we investigate two axes of behavioral plasticity that may impact a focal species’ participation in mixed flocks: ecological selectivity and flexibility. Flexibility of a species reflects the degree to which birds can adjust their foraging behavior to match or avoid the other flock members’ niche, and selectivity refers to the probability that a species will join a flock based on the ecological niche its members are exploiting. Selectivity refers to birds’ tendency to participate in a subset of flocks found in the community based on flock mate characteristics that maximize the benefits they may obtain. Selective species may prefer flock mates with different niches than themselves, prioritizing a reduction in competition or they may prefer birds of similar niches for the facilitative benefits they provide. Selectivity has been measured in classic mixed-species flock co-occurrence literature using static species-level traits such as body size and fixed foraging guild metrics. Meanwhile, looking at flexibility allows us to account for how species may adjust their behavior to maximize flocking benefits. Flexible species may have the ability to forage successfully in a wider variety of flock compositions and to tailor their foraging behavior to the composition of each flock they join. Furthermore, this behavioral plasticity may permit flexible species to expand their niche beyond the strata in which they may normally forage with conspecifics or alone.

When a focal species exhibits ecological flexibility, we expect to see negative or positive correlations between their flocking behavior and that of their flock mates. A negative correlation indicates flexible divergence in foraging niche (Figure 1A), while a positive correlation indicates flexible matching in foraging niche (Figure 1C). It is important to note that flexible matching does not mean species’ overlap with their flock mates’ behavior exactly, but that their behavior moves towards that of their flock mates. Species that are not flexible will not show significant changes in their foraging behavior as they participate in flocks with varied flock mate foraging behavior (Figure 1B). Meanwhile, selective species will join only a subset of the flocks seen in their environment. Those that selectively avoid competition with ecologically similar flock mates may only be seen in flocks where flock mates occupy dissimilar niches (Figure 1D), and those that selectively seek facilitative interactions may only be seen in flocks where flock mates occupy similar niches (Figure 1F). Species may also exhibit both flexibility and selectivity by actively converging upon or avoiding flock mates via changes in behavior within the subset of flocks they selectively participate in.

**Figure 1.**
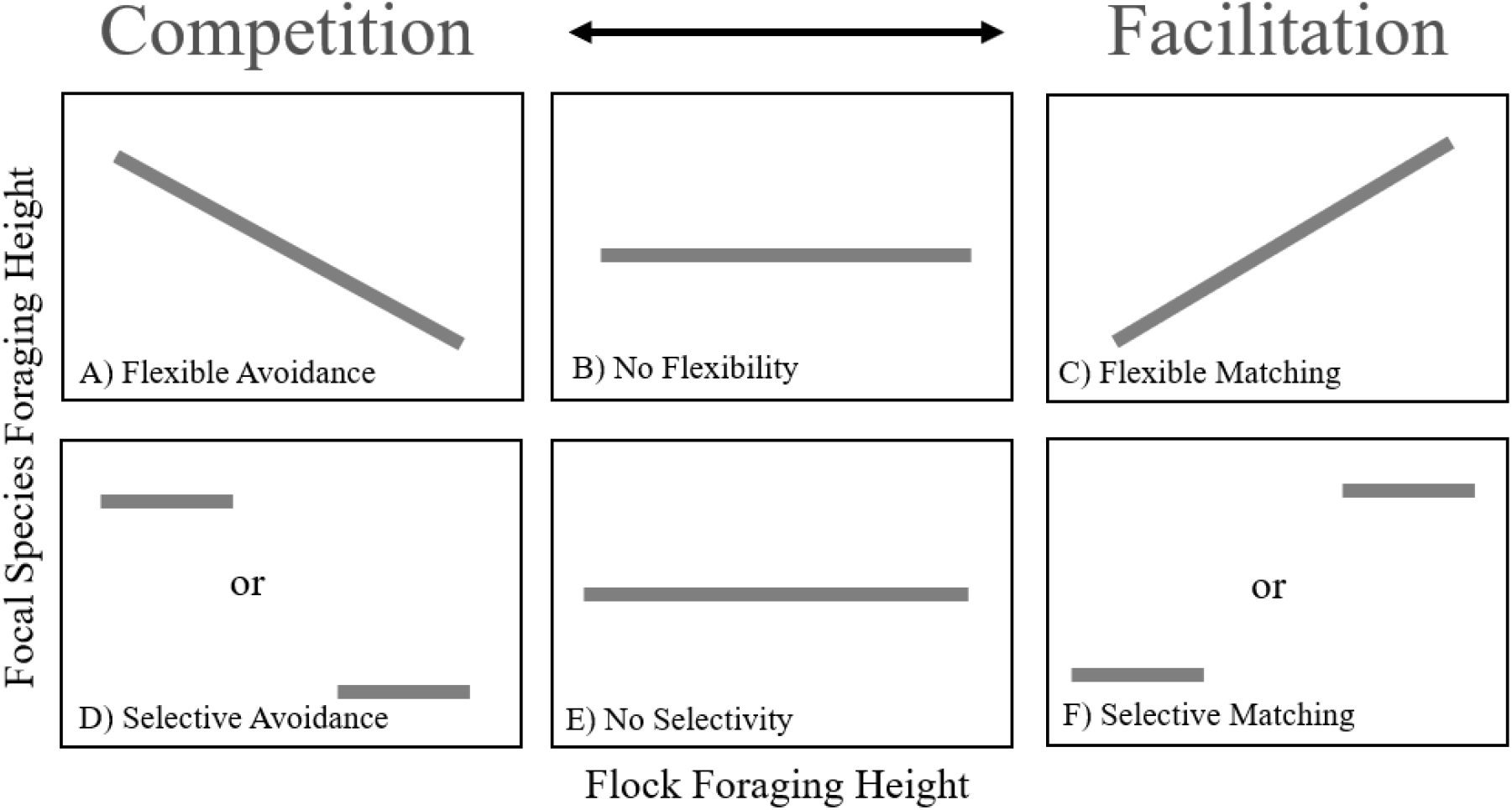
Hypothetical patterns of selective and flexible behaviors of a focal species and its flock mates. Foraging height is used for clarity, though any continuous metric of niche space could be used in its stead. Species may either avoid overlapping in foraging height with their flock mates by flexibly shifting their foraging height away from their flock mates (A) or selectively participating in flocks of different heights either lower or higher than the focal species behavior (D). Alternatively, species may match their flock mates’ foraging height via flexible changes in behavior (C) or by selectively flocking with flocks of similar heights, matching at either high or low foraging heights(F). Species that don’t exhibit flexible behaviors will show neither a negative (A) or positive (C) correlation with their flock mates’ behavior (B). Non-selective species will participate in flocks containing the full range of behaviors found in the system (E).

In addition to looking at how species’ patterns of participation correspond with the ecological niche of flocks, this framework may also be used to investigate relationships between pairs of closely interacting species within flocks. These high levels of interaction may arise for a number of reasons - sharing of territories, flushing of prey, particularly relevant information transfer (Seppänen et. al. 2007; Gyimesi et al. 2012; Johnson et al. 2018) - and may be mutualistic or commensalistic. In our study system, previous studies have shown a particularly tight relationship between two flocking species - the purple-backed fairywren (*Malurus assimilis)* and the splendid fairywren (*Malurus splendens)*. At our site these two fairywrens species show high rates of territory overlap, territory co-defense and group foraging behaviors. When closely associated with splendid fairywrens, purple-backed fairywrens were less vigilant, spent more time foraging and showed significant fitness benefits (Johnson et al. 2018). Due to the nature of their relationship, we predict that the flexibility of behavior shown by purple-backed and splendid fairywrens will be more tightly linked with one another’s foraging behavior than that of the flocks they participate in.

In this study we explore how species’ foraging flexibility and selectivity allow them to match or avoid similarly foraging flock mates within mixed-species flocks in South Australia. We examine patterns of flexibility and selectivity in flock participation via the following methods. 1) Using foraging stratum as a proxy for niche space, we map a species’ observed flocking stratum against that of its flock mates to examine patterns of selectivity and flexibility across species in the flocking community. 2) We assess patterns of foraging flexibility in a broader context via comparison of species’ foraging behavior in and out of mixed-species flocks. 3) We investigate selectivity and flexibility in the context of two species--purple-backed and splendid fairywrens--which have been previously shown to interact closely. Prior work shows that these two species co-defend territories, and association with splendid fairywrens confers foraging and predator vigilance benefits to purple-backed fairywrens (Johnson et al. 2018). By exploring these patterns of selectivity and flexibility in and out of mixed-species foraging flocks, we gain a deeper understanding of how different species navigate the interplay between competitive and facilitative behaviors in mixed-species groups.

## Methods

### Field Site

We conducted this work at Brookfield Conservation Park (−34.35°, 139.50°) in the lower Murray-Darling River Basin in South Australia from August 2019 to December 2019. Our season began just before the beginning of the austral spring and bird breeding season, continuing through the end of the breeding season when most species had fledged young. The landscape at the site is predominantly covered by second-growth mallee eucalypt woodland and chenopod scrub (Figure 2). This open landscape allows for detailed behavioral observations of mixed-species flocks. Mixed-species flocks at this site are ephemeral and dynamic, with flocks forming and dissolving throughout the day and composition changing frequently. The flocks consist primarily of insectivorous species, though nectarivores and frugivores like honeyeaters join as well.

**Figure 2.**
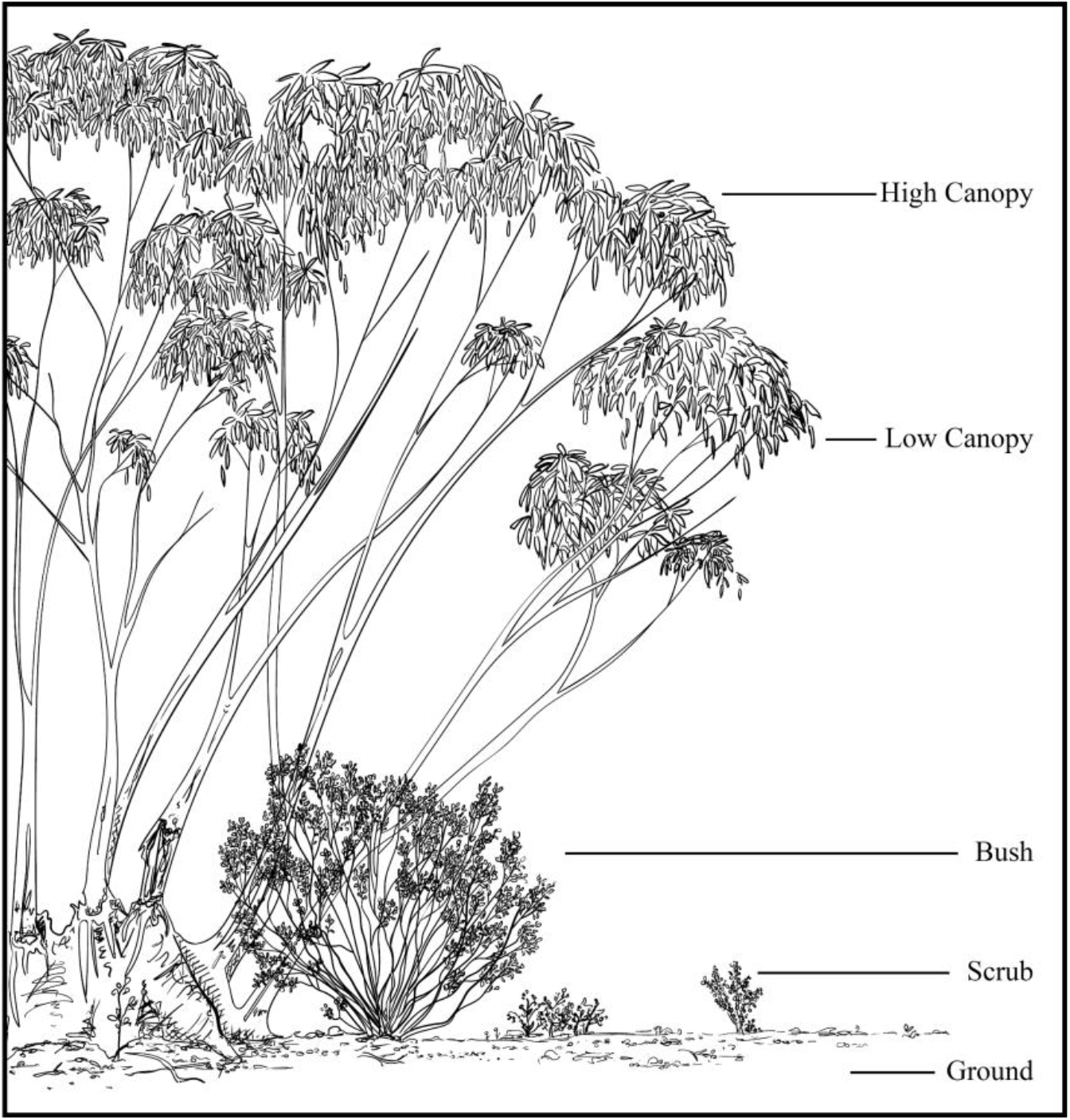
Observations of species’ flocking heights were split into five strata. 1) Ground – bare ground with no plant cover. 2) Scrub – in and around plants between 5cm and 20 cm in height. 3) Bush - in bushes or plants above 20 cm in height. 4) Low canopy – in the bottom half of a tree whether on open branches or in leaves. 5) High canopy – in the upper half of a tree, again either on open branches or in leaves. Illustration by AEJ.

#### Flock Observations

Throughout the season we opportunistically observed flocks of birds to determine species composition and ecological niche breadth of flock members. We recorded flock observations throughout the day, beginning at dawn and ending before dusk. We characterized flocks as mixed-species flocks if they contained two or more species and were either actively moving through the landscape in a coordinated fashion or seen together in one location for a period of more than five minutes. We note that many mixed-species flock studies do not include stationary flocks due to the potential influence of stationary resources (Greenberg 2000). However, since the presence of stationary resources such as flowering or fruiting trees was easily identifiable at our site, we included stationary flocks in our analysis except when a flowering or fruiting tree was the obvious cause of an aggregation. For each flock observed we recorded the location, species composition, number of individuals of each species, and range of strata in which each species was seen.

#### Foraging Strata

For every species in each flock, we recorded the highest and lowest foraging strata in which its members were observed. We designated foraging strata by separating the environment into five distinct, ecologically relevant categories – ground, scrub, bush, low canopy, and high canopy (Figure 2). The descriptions used to designate each category are as follows: 1) ground was defined as open ground, free of vegetation, 2) scrub consisted of plants taller than 5 cm but shorter than 20 cm—typically saltbush (*Atriplex* spp.), 3) bushes varied in height but the dominant species at our site were sheepbush (*Geijera linearifolia*), senna (*Senna artemisioides* ssp.), and bluebush (*Maireana* spp.), all of which are easily distinguishable from trees by their shape and leaves, 4) low canopy consisted of the lower half of branches and leaves on an individual tree (primarily *Eucalyptus* spp. and *Myoporum platycarpum*) (5) and high canopy consisted of the upper half of branches and leaves on an individual tree.

#### Individual Foraging Observations

To explore how joining mixed-species flocks influences the ecological niches of focal species, we took opportunistic foraging observations of birds seen both in and out of flocks for a select set of focal species. For each focal species individual seen we observed one distinct foraging event (a singular attempt to capture prey, successful or not) during which we recorded whether or not the individual was participating in a mixed-species flock, the stratum in which the foraging event took place, the cover category of the location of the event and a GPS point. The stratum designations used in the environment are the same as described in flocking observations above.

##### Analysis

For each flock observation, we filtered out all observations of species for which we had not recorded complete stratum data, i.e., both the highest and lowest strata for which a species was observed in each flock. We completed the following analyses on all species that were observed in at least 20 flocks over the course of the field season.

#### Focal Species and Flock Mate Height Midpoints

We assigned each stratum to an ordinal bin, using integer values 1 through 5 representing ground, scrub, bush, low canopy, and high canopy respectively. For each species in a flock, we calculated the midpoint of its foraging height by averaging the ordinal values of the lowest and highest strata in which members of that species were seen in that flock (Bangal et al. 2021). We also calculated the average midpoint of each focal species’ flock mates separately for every flock.

#### Flocking Flexibility Analysis

To identify whether species demonstrated flexibility in foraging height during flock participation, we tested for associations between the median foraging heights of each focal species and the average median heights of their flock mates using simple (bivariate) linear regressions with normally-distributed errors. Because the response data are not strictly continuous, we also analyzed the data using ordered logistic regression models (see Supplemental Materials). However, as the patterns we observed when using that approach did not differ materially from those found using simple linear regressions, we report the results of the former approach here for ease of interpretation.

To test for the effect of specific flock mates on behavior of a focal species, we ran a separate linear regression to highlight the interactions between two species known to form tight heterospecific associations during the breeding season—the purple-backed and splendid fairywrens (Johnson et al. 2018). In this model, we compared the midpoint stratum of purple-backed fairywrens with that of splendid fairywrens to determine whether the two species demonstrated flexible matching behavior with one another rather than flexibly responding to the average height of their flock mates.

#### Selectivity Analysis

We determined whether species selectively participated in flocks based on the flock’s height using permutation tests. For each focal species and every flock, we first calculated the average of all other species’ median heights in the flock, including flocks in which the focal species was not a participant. From this distribution of flock heights, we drew 10,000 random samples without replacement, with each sample equal in size to the number of flocks in which the focal species was seen. At each draw, we compared the empirical distribution of flock heights for the focal species to the resampled distribution using a two-sided *t*-test. We use the resulting distribution of *t* values from these tests as an indication of species selectivity: positive *t* statistics reflect species that (on average) selected flocks with higher median flock heights than expected by chance, whereas negative *t* statistics reflect those selecting flocks with lower median flock heights than expected by chance.

#### Conspecific and Mixed-species Flock Foraging Strata

We compared species’ foraging heights when in mixed-species flocks to that when foraging with conspecifics only. For each species that we observed foraging at least 15 times both in and out of mixed-species flocks, we used a Fisher’s exact test to determine whether the frequency of foraging across strata was similar in the two flock types.

## Results

### Flock Attributes

Over the course of the field season, we observed 306 independent mixed-species flocks made up from a pool of 38 flocking species. The average flock richness was 4.20 (sd+/-1.97) species; the most species rich flock observed was made up of 10 species.

### Flexibility and selectivity

All 13 species analyzed had positive or neutral relationships between their height midpoints and the average height midpoints of their flock mates (Figure 3). Of these 13 species, 8 showed patterns of flexible matching, having significantly positive correlations with the average flocking height of their flock mates, zero species showed patterns of flexible avoidance or a negative correlation with the average flocking height, and 5 species did not show a significant correlation (Table 1).

**Figure 3.**
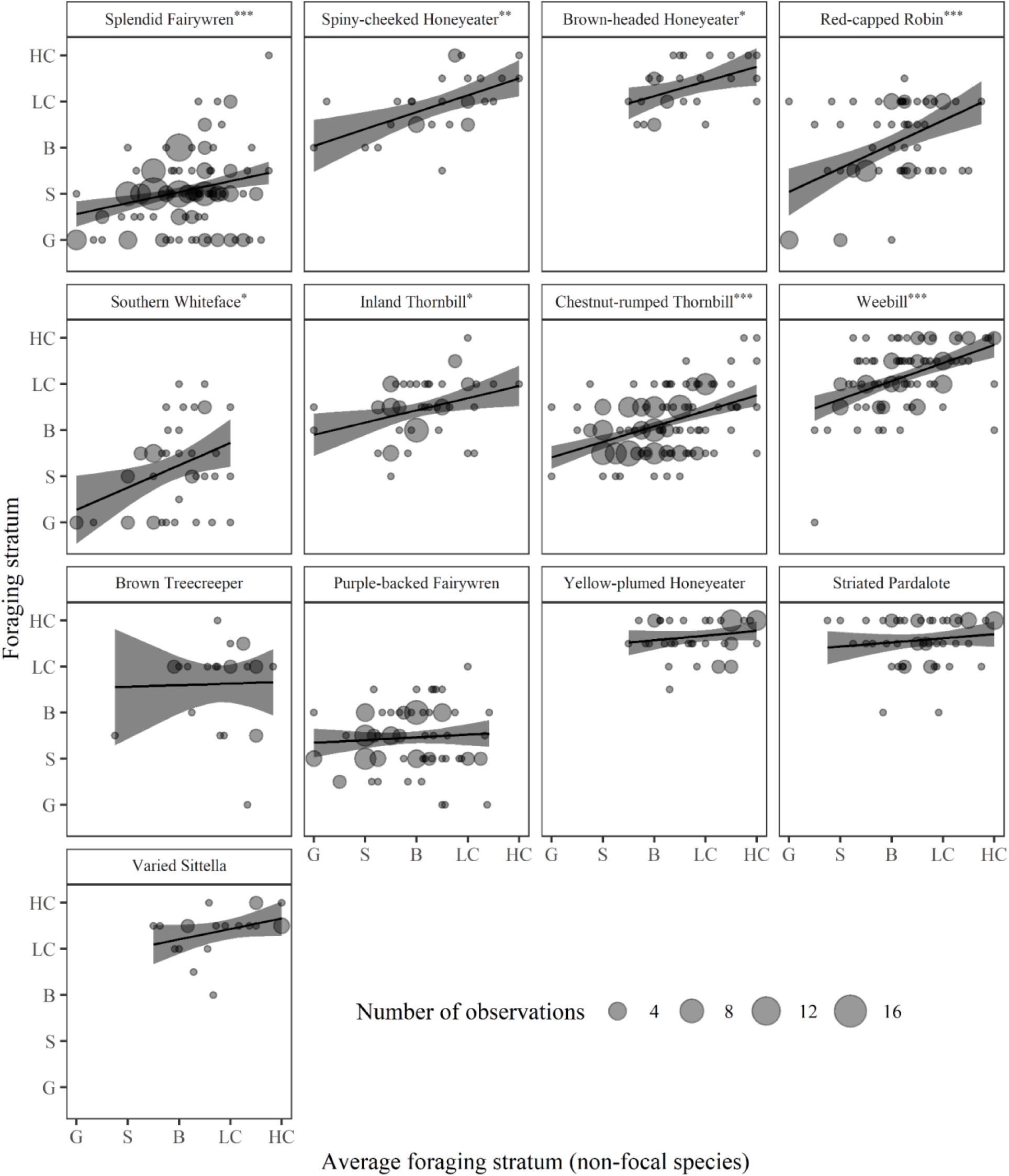
Relationship between a focal species foraging height midpoint and the average of its flock mates’ height midpoints. The size of the points indicates the number of flock observations that fall in the same place on the graph. A * indicates p<0.05, ** indicates p<0.01, and *** indicates p<0.001.

**Table 1.** Species Flocking Stratum, Flexibility and Selectivity. Columns marked with a * indicate p<0.05, ** indicates p<0.01, *** indicates p<0.001, a + in the selectivity column indicates that a species selectively participated in flocks with higher average stratums than expected by chance, while a - means a species selectively participated in flocks with a lower average stratum than expected by chance, and --- indicates insufficient data to run an analysis.

|  |  | Average<br>strata<br>midpoint | Within flock<br>flexibility | Selectivity | Conspecific vs.<br>mixed-species<br>flock flexibility |
| --- | --- | --- | --- | --- | --- |
| *Acanthagenys*<br>*rufogularis* | Spiny-cheeked<br>Honeyeater | 3.93 ± 0.65 | ** | + | --- |
| *Acanthiza apicalis* | Inland Thornbill | 3.46 ± 0.60 | * | Not selective | Not flexible |
| *Acanthiza*<br>*uropygialis* | Chestnut-rumped<br>Thornbill | 3.11 ± 0.63 | *** | Not selective | * |
| *Aphelocephala*<br>*leucopsis* | Southern Whiteface | 2.16 ± 0.94 | * | - | Not flexible |
| *Climacteris*<br>*picumnus* | Brown Treecreeper | 3.63 ± 0.92 | Not flexible | + | --- |
| *Daphoenositta*<br>*chrysoptera* | Varied Sittella | 4.40 ± 0.49 | Not flexible | + | Not flexible |
| *Lichenostomus*<br>*ornatus* | Yellow-plumed<br>Honeyeater | 4.67 ± 0.40 | Not flexible | + | --- |
| *Malurus assimilis* | Purple-backed Fairywren | 2.46 ± 0.61 | Not flexible | - | ** |
| *Malurus splendens* | Splendid Fairywren | 2.04 ± 0.70 | *** | - | Not flexible |
| *Melithreptus*<br>*brevirostris* | Brown-headed<br>Honeyeater | 4.31 ± 0.54 | * | + | Not flexible |
| *Pardalotus*<br>*striatus* | Striated Pardalote | 4.58 ± 0.50 | Not flexible | + | Not flexible |
| *Petroica*<br>*goodenovii* | Red-capped Robin | 3.05 ± 0.95 | *** | Not selective | --- |
| *Smicrornis*<br>*brevirostris* | Weebill | 4.18 ± 0.67 | *** | + | *** |

Ten of 13 species showed selectivity (Figure 4, Table 1). Three species did not, having patterns of flock participation indistinguishable from random selection of flocks observed in the environment. Species’ selectivity appears to occur in the direction of their flocking preferences; species who’s average flocking midpoint was above 3 (the middle strata of bush) were seen in flocks that were at a higher average stratum than random, while species whose average flocking midpoint was lower than 3 were seen in flocks at a lower average strata than expected by chance (Table 1). Six of the 13 species showed patterns of both flexibility and selectivity (Table 1).

**Figure 4.**
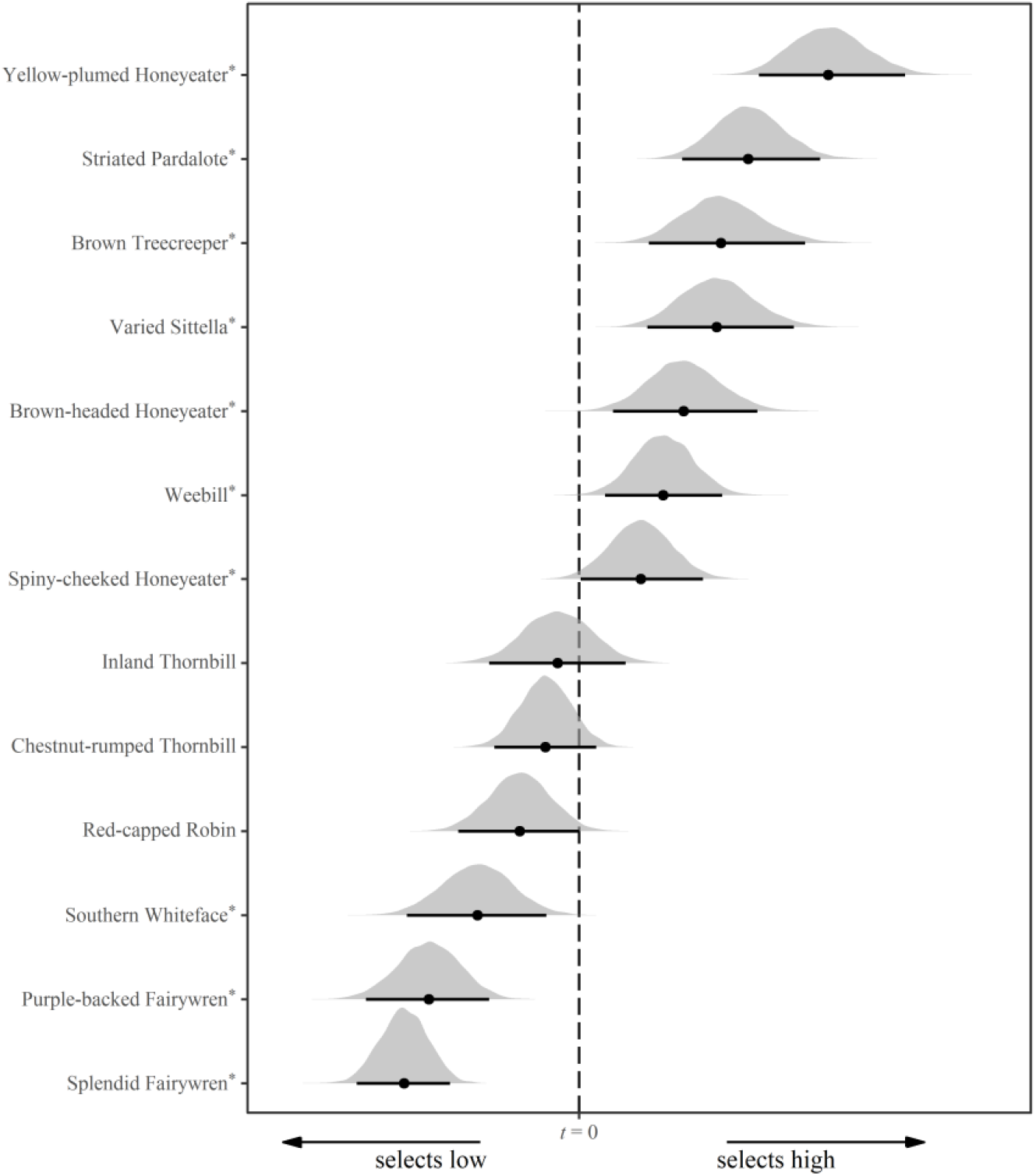
Representation of the 95% confidence intervals of 1000 randomizations of flock height for each species. Species for which the confidence interval overlaps zero do not participate in a subset of flocks that have significantly different average heights than the flocks observed. Species whose confidence intervals are to the right of 0 participate in flock that are significantly higher than the expected by chance, while species whose confidence intervals are to the left of 0 participated in flocks that are significantly lower in strata than expected by chance. All species that participate in subset of flocks significantly different than random are marked by *.

### Conspecific and Mixed Flock Foraging Strata

Three out of nine species had significantly different proportions of foraging strata observations when observed in mixed-species flocks than when foraging solo or with conspecifics (Figure 5). Two of the three species had a midpoint height that was strongly correlated with their average mixed flock mates’ height while the third’s, the purple-backed fairywren, flocking height was not significantly correlated with the average midpoint of its mixed flock mates (Table 1).

**Figure 5.**
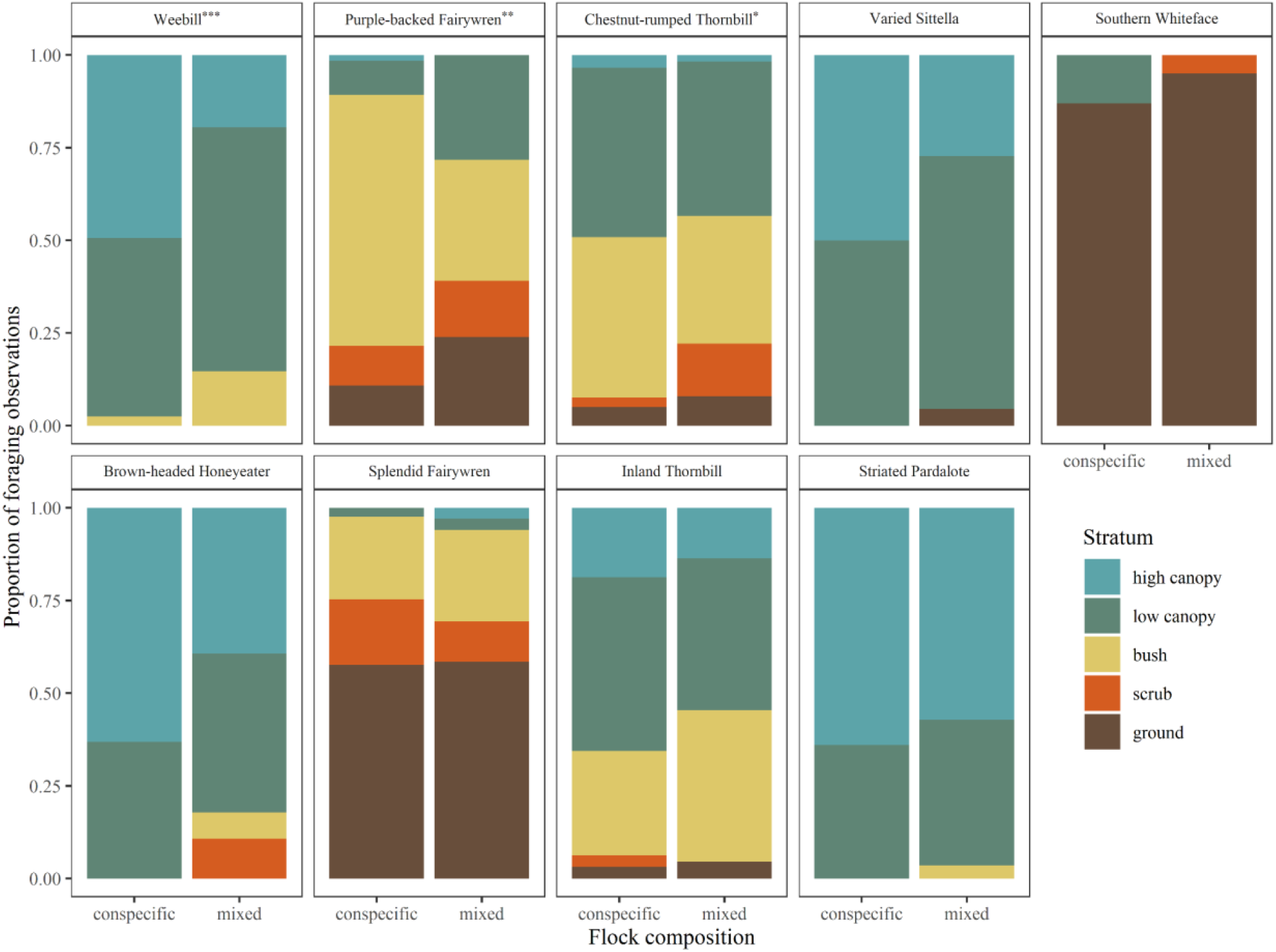
Proportion of observed foraging heights of flocking species in and out of flocks. Each color represents different foraging strata from ground to high canopy. Species who showed significant differences in the proportions of observations in each strata are marked with * for p<0.05, ** for p<0.01, and *** for p<0.001.

**Figure 6.**
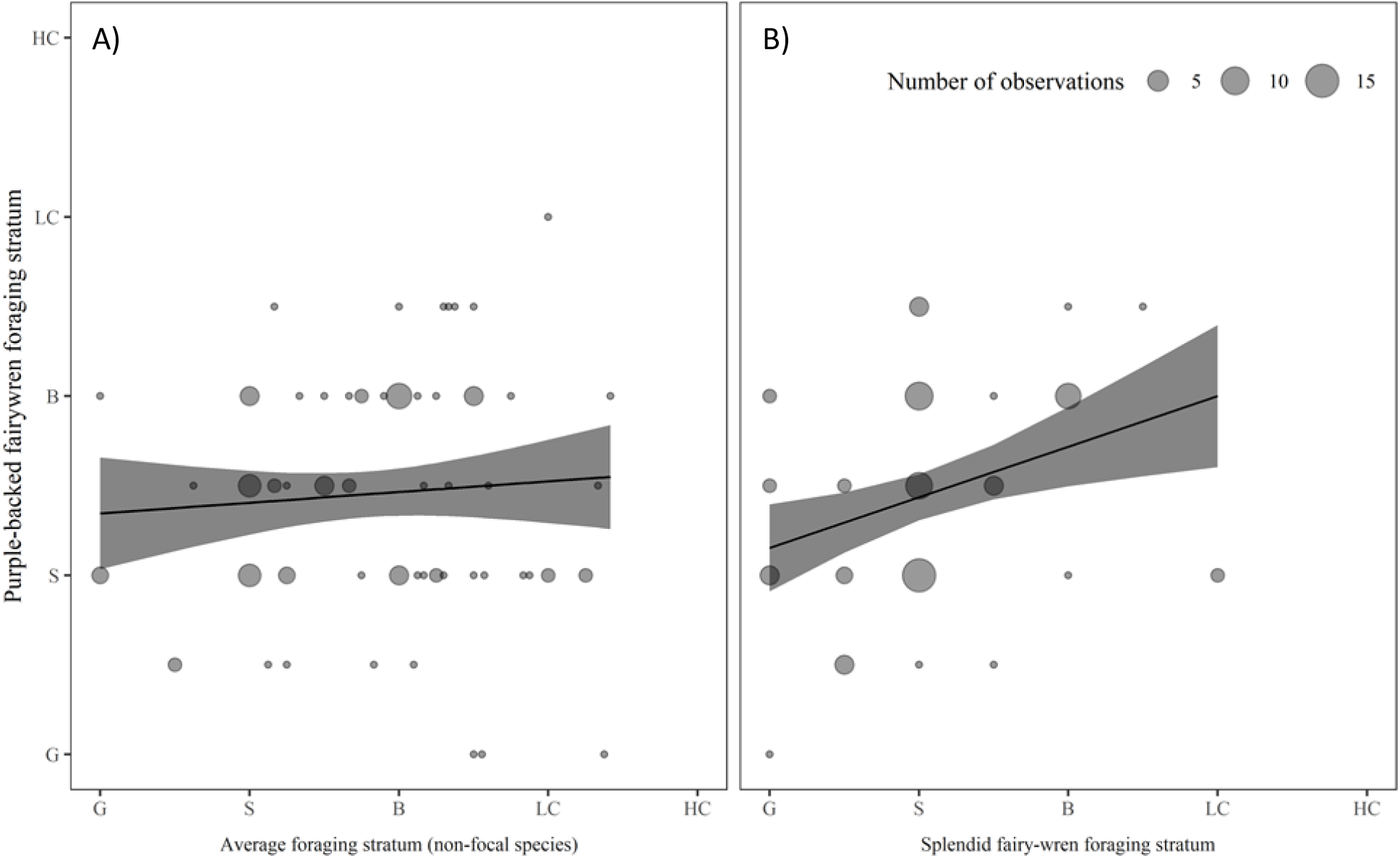
(A) Relationship between purple-backed fairywrens’ midpoint height and that of their average flock mates’ (R^2 = 0.00491, F(1,90)=0.44, p=0.51). (B) Relationship between purple-backed fairywrens height midpoints and the height midpoints of splendid fairywrens in the same flocks. The size of the points represents the number of flocks showing the same relationship (R^2 = 0.1021, F(1, 73) = 8.3, p = 0.005).

### Fairywren Case Study

Purple-backed fairywrens and splendid fairywrens were observed to co-occur in 75 flocks together. Purple-backed fairy wrens were observed participating in 92 mixed-species flocks, 81.52% of which contained splendid fairywrens. Splendid fairywrens were observed in 180 mixed-species flocks, 41.67% of which contained purple-backed fairywrens Though purple-backed fairywrens did not show a significant correlation in their midpoint height when compared with that of their flock mates (R^2 = 0.00491, F(1,90)=0.44, p=0.51), purple-back fairywrens and splendid fairywrens had significantly correlated height midpoints when they were seen in flocks together. (R^2 = 0.1021, F(1, 73) = 8.3, p = 0.005) (Figure 5). This suggests that purple-backed fairywrens may exhibit ecological flexibility with respect to the foraging strata of splendid fairywren flock mates but not the average foraging strata of other flock mates.

## Discussion

In this study, we explored how behavioral plasticity may shape mixed-species flocks by allowing birds to manage the effects of competition and facilitation with other flock members. Species that join mixed-species flocks in order to gain benefits of sharing ecological niche space with heterospecific individuals can maximize these benefits in two ways: by changing their foraging niche to match that of their flock mates (behavioral flexibility) or by selectively joining mixed-species flocks when they occur within their preferred foraging niche (selectivity). In this study, we find examples of species employing both strategies. Eight species showed patterns of flexible matching with their flock mates’ foraging strata, 10 species showed selectivity for flocks that corresponded with their preferred foraging stratum, (e.g., canopy species selected for canopy flocks), and five species employed a combination of the two strategies. Notably, none of the 13 species showed patterns of competitive avoidance either through selective participation in flocks with differently foraging species or by adjusting behavior to avoid flock mates. Taken as a whole, this research supports the hypothesis that facilitation between ecologically similar species plays a key role in the formation and composition of mixed-species flocks. However, this data also shows that different species match the behavior of their flock mates via different methods, by flexibly converging upon their flock mates’ height, selectively picking flocks of certain foraging heights, or a combination of the two.

Some species depended solely on behavioral flexibility rather than selectivity to manage their species interactions. In our analysis, three species showed patterns of flexible matching without showing selective preferences for the height of the flocks they participated in. Two of these species, chestnut-rumped thornbills (*Acanthiza uropygialis*) and weebills (*Smicrornis breivirostris*), were also found to exhibit behavioral flexibility outside of mixed-species flocks, foraging in significantly different strata when solo or with conspecifics than when foraging in mixed-species flocks. Taken together, these two points of evidence imply that these species actively change their behavior to remain close to their flock mates, and in doing so expand their foraging niche beyond the strata in which they normally forage. A similar result was found in a study by Farine and Milburn (2013) in buff-rumped thornbills that expanded their foraging strata in either direction, lower or higher, depending on the identity of thief flock mates. Many studies on behavioral niche expansion within mixed-species flocks have found that species take advantage of insects flushed by the movement of the flock through the environment (Munn 1986; Sridhar and Shanker 2014) with evidence for species switching from frugivory to insectivory as they join flocks (Valburg1992) and suggestions that behavioral matching occurs because species must be close to flock mates in order to benefit from the flushing of insects (Sridhar and Shanker 2014). Neither chestnut-rumped thornbills nor weebills typically catch insects on the wing (Vander Meiden, personal observations), suggesting that they may expand their niche in other ways when joining mixed-species flocks.

Other flocking species manage their interspecific interactions not via flexibility, but by selectively participating only in flocks with birds of similar foraging behaviors. In our system, ten species preferentially participated in flocks in which the average flocking height was significantly different than expected by chance in this environment. In each of these cases the non-random selection of flock mate height corresponded with the preferred strata of the focal species. Species whose average flocking midpoint was in the canopy – above a 3 – joined flocks that were significantly higher than the height of a random sampling of flocks, and species whose average flocking midpoint was in the understory – below a 3 – joined flocks that were significantly lower than random (Table 1). This pattern of behavior in which a species selectively engages with similarly flocking species could generate the type of trait assortment that has been found in other mixed-species flock systems (Sridhar et al. 2012; Bangal et al. 2021). Studies have shown that birds of similar foraging guilds, body size, and phylogenetic relatedness are more likely to occur in flocks together (Sridhar et al. 2012). Three of these ten selective species showed no pattern of flexible behavior in conjunction with their selective behavior (Table 1). This lack of flexible behavior once in flocks, may imply that these species join flocks for benefits that do not require more than general proximity to their flock mates, such as access to alarm calls warning of predator presence. In such cases there may be no additional benefit to tailoring behavior to that of the other species present. Conversely it may be that solely selective species are participating in flocks based on variables that are correlated with their preferred strata rather than participating in flocks based on the average height of the flock itself.

Other species may have the ability to flexibly adjust their foraging behavior but, rather than matching the behavior of the whole flock may only do so in response to the presence or behavior of a particular species. Purple-backed fairywrens had significantly different foraging behavior when in mixed-species flocks than when outside of them, expanding their foraging into both higher and lower strata in flocks. However, purple-backed fairywrens did not show a pattern of flexibly matching flock height when participating in mixed-species flocks. The purple-back fairy wrens did, however, show a significant correlation in flocking strata with that of splendid fairywren flock mates. Previous research at this site has shown that purple-backed fairy wrens consistently forage with splendid fairywrens and gain vigilance and foraging benefits by doing so (Johnson et al. 2018). Eighty-one percent of the mixed-species flocks where purple-backed fairywrens participated also contained splendid fairywrens. Meanwhile, most of the time (68.33%) splendid fairywrens were observed, purple-backed fairywrens were not present. Thus, it may be that purple-backed fairywrens, though not splendid fairywrens, primarily obtain flocking benefits via proximity to splendid fairywrens rather than the flock as whole. Thus, the proximity of other flock mates may be inconsequential. The strong relationship between the two fairywren species highlights the potential importance of dyadic relationships on patterns of selectivity and flexibility.

By looking at relationships between species and their behavior at the dyad level, we have the potential to gain greater clarity about the function and importance of species roles in mixed-species flocks. Studies have shown that some nuclear species provide high levels of vigilance and consistent alarm calls when predators are near, allowing flock mates to increase their foraging rate or expand their foraging niche (Dolby and Grubb 1998). Further, meta-analyses have shown flocking benefits are typically received by satellite species not nuclear species (Sridhar et al 2009). Therefore, when looking at behavioral flexibility, we would expect multiple flocking species’ behavior to show strong correlations with the behavior of the nuclear species. In cases where flocking benefits are primarily provided via association with nuclear species and not via the flock, we would expect behavioral correlations between satellite and nuclear species to be stronger than those between satellite species and other flock mates. By looking at a species’ flexibility and selectivity at both the flock and dyadic level we can learn more about the relative importance of different species on flock benefits such as niche expansion and have a way of numerically identifying nuclear species.

Looking at species’ responses to the behavior of flock mates via patterns of selectivity and behavioral flexibility can provide key insights into which species we expect to interact in flocks and what those interactions are like respectively. Crucially, our results show that different species use different strategies to modulate their behavior in response to that of flock mates. Thus, analyses of flock mate co-occurrence and interactions using static species-level traits are incomplete without accounting for the potential behavioral flexibility of flock mates. By looking at where each species falls on the axes of flock selectivity and behavioral flexibility, we can begin to infer the types of benefits received by participation. Use of this selectivity/flexibility framework across additional niche parameters can help us narrow down how flocking benefits are received and which aspects of a species’ niche are constrained or expanded by competitive or facilitative interactions with flock mates. Finally, use of this analysis in conjunction with social network analysis could move the literature past rough nuclear/satellite species designations by looking at the degree to which dyadic relationships between nuclear and satellite species influence species’ participation and behavior in mixed-species flocks and comparing that to how satellite species respond to the presence and behavior of other flocking species or the flock as a whole.

## Notes

### Competing Interest Statement

The authors have declared no competing interest.

